## Supplementary file for "Characterization of two non-competing antibodies to influenza H3N2 hemagglutinin stem reveals its evolving antigenicity"

**Table S1. Cryo-EM data collection, refinement, and validation statistics.**

| | H3N8 HA + AG2-G02 (EMD-48873) (PDB 9N4E) | H3N8 HA + 2F02 (EMD-48874) (PDB 9N4F) |
|---|---|---|
| **Data collection and processing** | | |
| Magnification | 81,000 | 81,000 |
| Voltage (kV) | 300 | 300 |
| Electron exposure ($e^{-}/Å^2$) | 57.35 | 57.35 |
| Defocus range (μm) | -0.5 to -3.0 | -0.5 to -3.0 |
| Pixel size (Å) | 0.53 | 0.53 |
| Symmetry imposed | C1 | C1 |
| Initial particle images (no.) | 552,543 | 459,706 |
| Final particle images (no.) | 326,211 | 120,986 |
| Map resolution (Å) | 2.6 | 2.71 |
| FSC threshold 0.143 | | |
| Map postprocessing | DeepEMhancer | DeepEMhancer |
| **Refinement** | | |
| Initial model used (PDB code) | N/A | N/A |
| Model resolution (Å) | | |
| FSC threshold | | |
| Model resolution range (Å) | | |
| Map sharpening *B* factor ($Å^2$) | N/A | N/A |
| Model composition | | |
| Non-hydrogen atoms | 16,662 | 16,473 |
| Protein residues | 2,124 | 2,124 |
| Ligands | | |
| *B* factors ($Å^2$) | | |
| Protein | | |
| Ligand | | |
| R.m.s. deviations | | |
| Bond lengths (Å) | 0.005 | 0.006 |
| Bond angles (°) | 0.980 | 1.021 |
| Validation | | |
| MolProbity score | 1.79 | 2.18 |
| Clashscore | 7.35 | 10.92 |
| Poor rotamers (%) | 1.49 | 3.67 |
| Ramachandran plot | | |
| Favored (%) | 96.18 | 96.85 |
| Allowed (%) | 3.82 | 3.15 |
| Disallowed (%) | 0.00 | 0.00 |

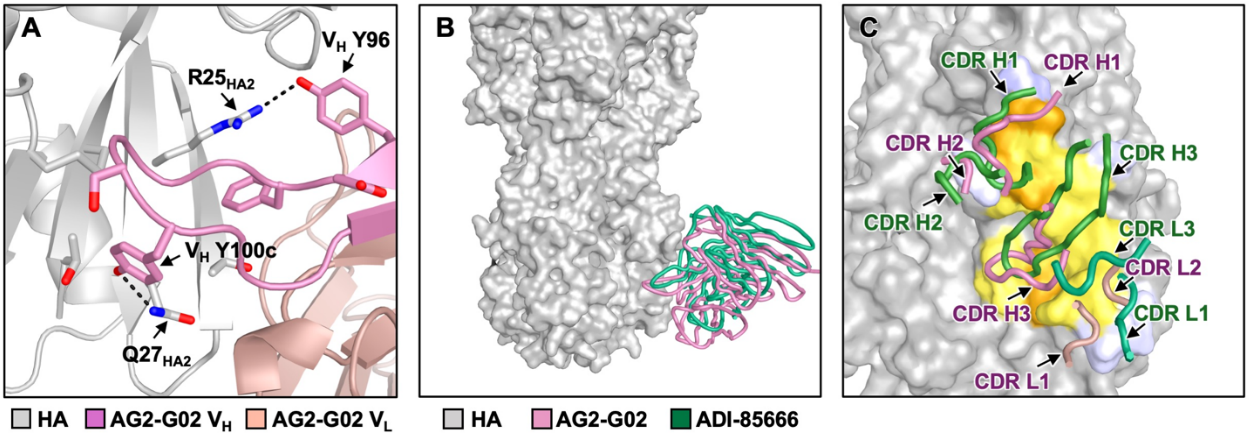


**Figure S1. Binding of AG2-G02 in comparison to other stem-targeting antibodies.**

**(A)** Interaction between the CDR H3 loop of AG2-G02 and the basal β-sheet that is formed by both HA1 and HA2 subunits in the HA stem. The side chains of key interacting residues are shown as sticks representation. Black dashed lines represent H-bonds. HA is depicted in gray, whereas the heavy and light chains of AG2-G02 are in pink and salmon, respectively.

**(B)** The binding of AG2-G02 (pink) and ADI-85666[1] (green, PDB 9BDF) to HA is compared.

**(C)** Interaction between the HA stem and CDRs of AG2-G02 as well as ADI-85666. Heavy and light chains of AG2-G02 are shown in pink and salmon, respectively. Heavy and light chain of ADI-85666 are in green and teal, respectively. HA1 and HA2 residues that are in the epitope of both antibodies are in orange and yellow, respectively. Epitope residues that are not shared by the two antibodies are in lavender.

**(B-C)** HA is shown as grey surface.

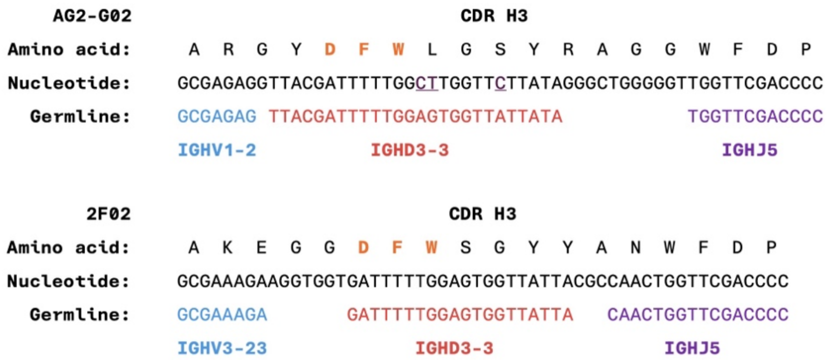


**Figure S2. Analysis of the CDR H3 sequences of AG2-G02 and 2F02.**
Germline analysis of the CDR H3 sequences of AG2-G02 and 2F02. Sequences corresponding to the proposed IGHV, IGHD, and IGHJ germline genes are highlighted in blue, red, and purple, respectively. Somatically mutated nucleotides are underlined. The spaces between the V-D and D-J junctions represent N-nucleotide additions. The "DFW" motif is highlighted in orange.

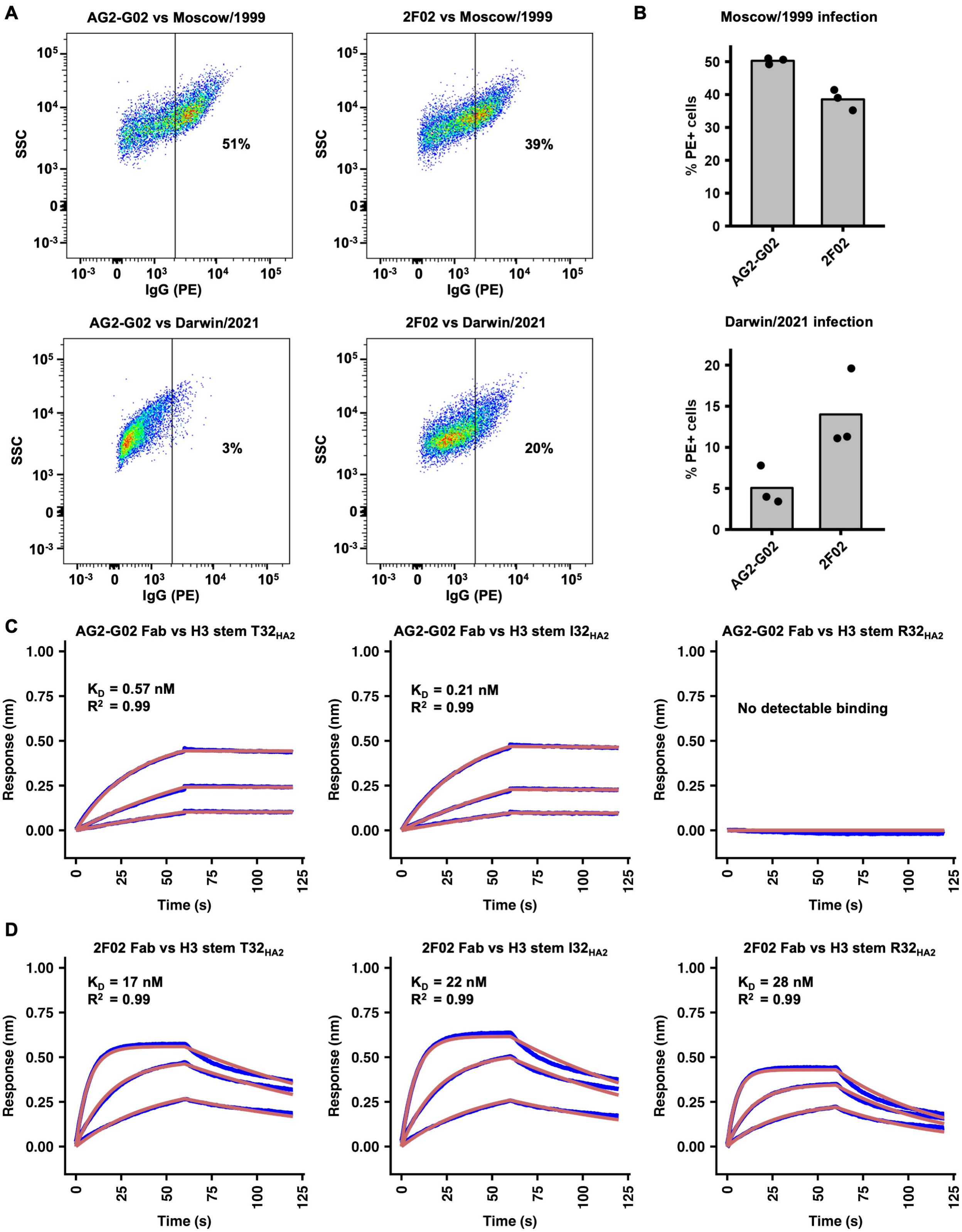


**Figure S3. Binding of AG2-G02 and 2F02 to infected cells and HA stem variants.**

**(A-B)** Binding of AG2-G02 IgG and 2F02 IgG to MDCK-SIAT1 cells infected with H3N2 A/Moscow/10/1999 (Moscow/1999) virus or H3N2 A/Darwin/6/2021 (Darwin/2021) virus at an

MOI of 0.1 was measured by flow cytometry. **(A)** Representative flow cytometry scatter plots from three biological replicates are shown. **(B)** The average frequency of PE-positive cells across three biological replicates is shown as bar plots. Each data point represents on replicate. **(C-D)** Biolayer interferometry was used to measure the binding kinetics of **(C)** AG2-G02 Fab and **(D)** 2F02 Fab against different variants of H3 stem, which was an HA stem construct designed based on H3N2 A/Finland/486/2004 HA[2], containing either Thr, Ile, or Arg at position 32 of the HA2. The Y-axis indicates the signal response. The blue lines depict the response curves, while the red lines represent the 1:1 binding model. Binding kinetics were assessed at three Fab concentrations: 300 nM, 100 nM, and 33 nM. The dissociation constants ($K_D$) and the goodness of fit values ($R^2$) are shown.

| Strain | HA2 positions 8 to 53 |
|---|---|
| **H3N8 Alberta/2017** | GFIENGWEGMIDGWYGFRHQNSEGTGQAADLKSTQAAIDQINGKLN |
| **H3N2 HK/1968** | GFIENGWEGMVDGWYGFRHQNYEGTGQAADLKSTQAAINQINGKLN |
| **H3N2 Philippines/1982** | GFIENGWEGMVDGWYGFRHQNSEGTGQAADLKSTQAAINQINGKLN |
| **H3N2 Beijing/1992** | GFIENGWEGMVDGWYGFRHQNSEGTGQAADLKSTQAAINQINGKLN |
| **H3N2 Wuhan/1995** | GFIENGWEGMVDGWYGFRHQNSEGTGQAADLKSTQAAINQINGKLN |
| **H3N2 Moscow/1999** | GFIENGWEGMVDGWYGFRHQNSEGTGQAADLKSTQAAINQINGKLN |
| **H3N2 NY/2004** | GFIENGWEGMVDGWYGFRHQNSEGIGQAADLKSTQAAINQINGKLN |
| **H3N2 Wisconsin/2005** | GFIENGWEGMVDGWYGFRHQNSEGIGQAADLKSTQAAINQINGKLN |
| **H3N2 Uruguay/2007** | GFIENGWEGMVDGWYGFRHQNSEGIGQAADLKSTQAAIDQINGKLN |
| **H3N2 Switzerland/2013** | GFIENGWEGMVDGWYGFRHQNSEGRGQAADLKSTQAAIDQINGKLN |
| **H3N2 Darwin/2021** | GFIENGWEGMMDGWYGFRHQNSEGRGQAADLKSTQAAIDQINGKLN |

**Figure S4. Sequence variation at HA2 position 32 of different H3 strains.**
The amino acid sequence at HA2 position 32 of an H3N8 and various H3N2 strains are shown, with threonine (T) shown in orange, isoleucine (I) in brown, and arginine (R) in green.

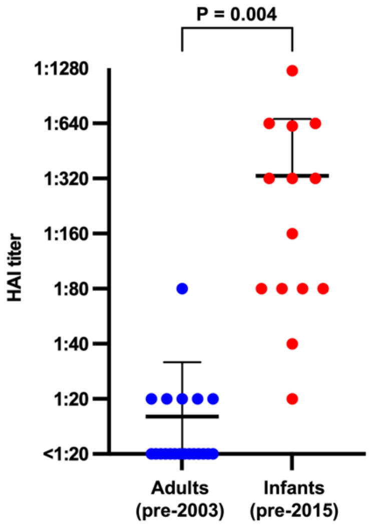


**Figure S5. Difference in cross-reactivity between plasma samples.**

Plasma samples from 15 infants under 2 years old and pre-2003 adults were analyzed by a hemagglutination inhibition (HAI) assay against the H3N2 A/Darwin/9/2021 virus. Each data point represents one plasma sample. Error bars represent standard deviations. The p-value was determined using a two-tailed Student's t-test.

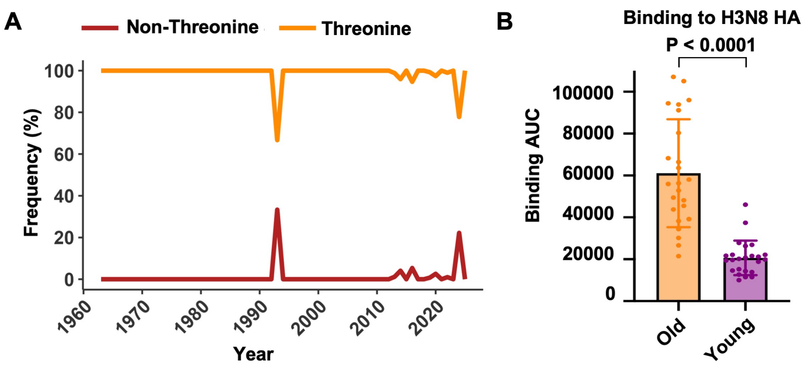


**Figure S6. Cross-reactivity of plasma samples to an avian H3 HA.**

**(A)** The percentage of occurrence of threonine at HA2 position 32 of avian H3 influenza virus was analyzed over the years. This analysis was based on 3,150 avian H3 HA sequences from the GSAID database[3]. The x-axis represents the years, while the y-axis shows the percentage of strains containing threonine (orange) or non-threonine (red) at HA2 position 32.

**(B)** ELISA showing the binding activity of plasma samples from 25 individuals over 60 years old (orange) and that from individuals aged 17-25 (purple) against avian H3N8 A/mallard/Alberta/362/2017 HA, which has $T32_{HA2}$. The y-axis represents the area under the curve (AUC) of four serial 10-fold dilutions of serum (1:100, 1:1,000, 1:10,000, and 1:100,000). The p-value was determined using a paired two-tailed Student's t-test. Error bars represent standard deviations.
